# Shifting resource limitation explains multiphasic patterns of density dependence

**DOI:** 10.64898/2026.07.16.738797

**Authors:** Andrew D. Letten, James A. Orr, Jan Engelstädter, Noelle A. Held, Christopher Klausmeier, Michael Manhart, Daniel B. Stouffer

## Abstract

Several recent studies have presented sublinear density dependence as a universal phenomenon across species, driven by factors unrelated to resource limitation (e.g. predation or non-resource based inhibition). Through a combination of bacterial growth experiments and mathematical modelling, we instead show that regimes of sublinear density dependence readily emerge under sequential shifts in resource limitation, without the need to invoke other processes. We nevertheless predict that superlinear density dependence, driven by standard resource limitation, will still predominate at both high and low densities.

## Main Text

Across the tree of life, per capita population growth typically declines as population density increases^1^. This negative feedback is a signature of diverse processes in ecology and evolution, from natural selection to ecosystem stability. Yet the shape of the relationship between growth and density remains contentious, and is thus a source of uncertainty in understanding and predicting population dynamics and ecosystem stability^2–8^.

On the one hand, mechanistic theory based on competition for limiting resources typically predicts the steepest declines in growth at high density (superlinearity)^2,9,10^. On the other hand, empirical evidence from experimental and observational time-series often points to more rapid declines in growth at low density (sublinearity)^4–6^. One proposed explanation for this apparent disconnect is that sublinearity in empirical data emerges primarily as a statistical artefact of regressing a variable upon itself^7,8^. Alternatively, it has recently been argued that processes unrelated to resource limitation account for observed patterns of sublinearity^4,5^. Here we show how resource limitation alone can explain recent observations of sublinear density dependence. Specifically, we compare simulations of consumer-resource models with high resolution bacterial growth assays to show how multiphasic patterns of super- and sublinear density dependence can emerge as different resources become limiting during population growth.

We first performed 24h batch culture assays with *E. coli* strain MG1655 in M9 minimal media at four initial glucose concentrations, with optical density (OD) measurements at three minute intervals. We quantified per capita growth rate by calculating the central difference of ln(OD) at each time point, which we then examined as a function of OD across each measurement interval (see Methods). At the lowest initial glucose concentration, we observed a strongly superlinear relationship between per capita growth rate and density (Fig. 1A). However, at higher initial glucose concentrations, the shape of density dependence was characterised by distinct regions of superlinearity at low and high density, joined by regions of sublinearity at intermediate densities (Fig. 1A & Fig. 2A). These inflection points were most conspicuous at the two highest glucose concentrations, particularly when per capita growth is averaged over multiple replicates (Fig. 2A). Inspection of the raw growth curves shows that the concave-up (i.e., sublinear) transitions, between regions of concave-down (i.e., superlinear) density dependence, correspond to comparatively abrupt downward shifts in growth rate. After each such transition, optical density continues to increase but at a conspicuously slower rate (Fig. 1B) until the population eventually saturates and per capita growth becomes zero.

**Figure 1:**
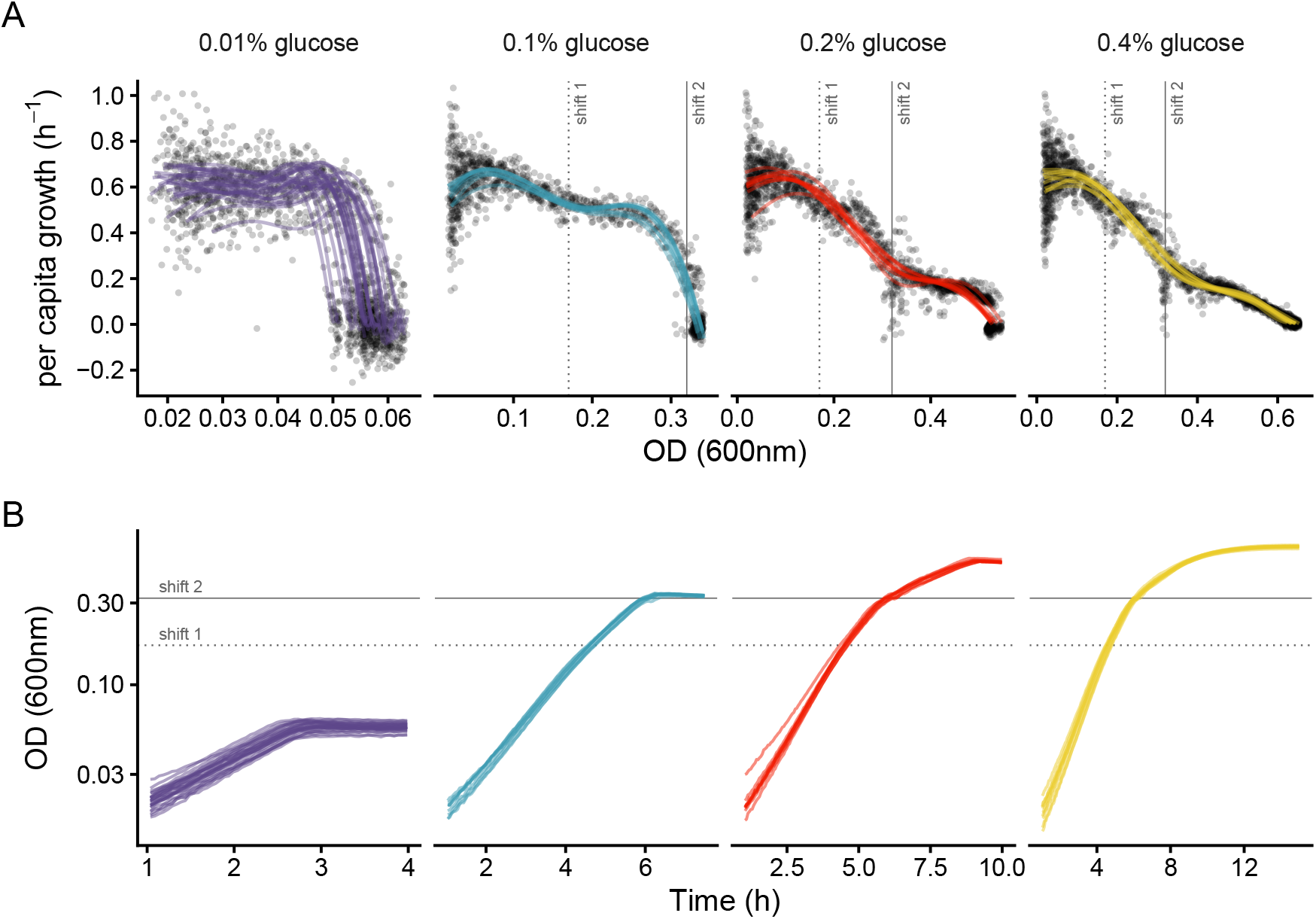
Observed *E. coli* population dynamics across different initial glucose concentrations. **(A)** Per capita growth rate (PGR) as a function of optical density. Points are estimates of PGR across each three minute time step across multiple replicates. Coloured lines are LOESS smoothers fitted to each replicate at each glucose concentration with a natural cubic spline with five degrees of freedom. **(B)** Raw optical density as a function of time (y-axis on a log scale) for each corresponding replicate in panel **A**. Vertical **(A)** and horizontal **(B)** dotted and solid lines denote discrete downward shifts in growth rate at consistent optical densities despite different initial glucose concentrations. Shift locations were estimated visually based on Fig. 2A (OD_1_ ≈ 0.17; OD_2_ ≈ 0.32).

**Figure 2:**
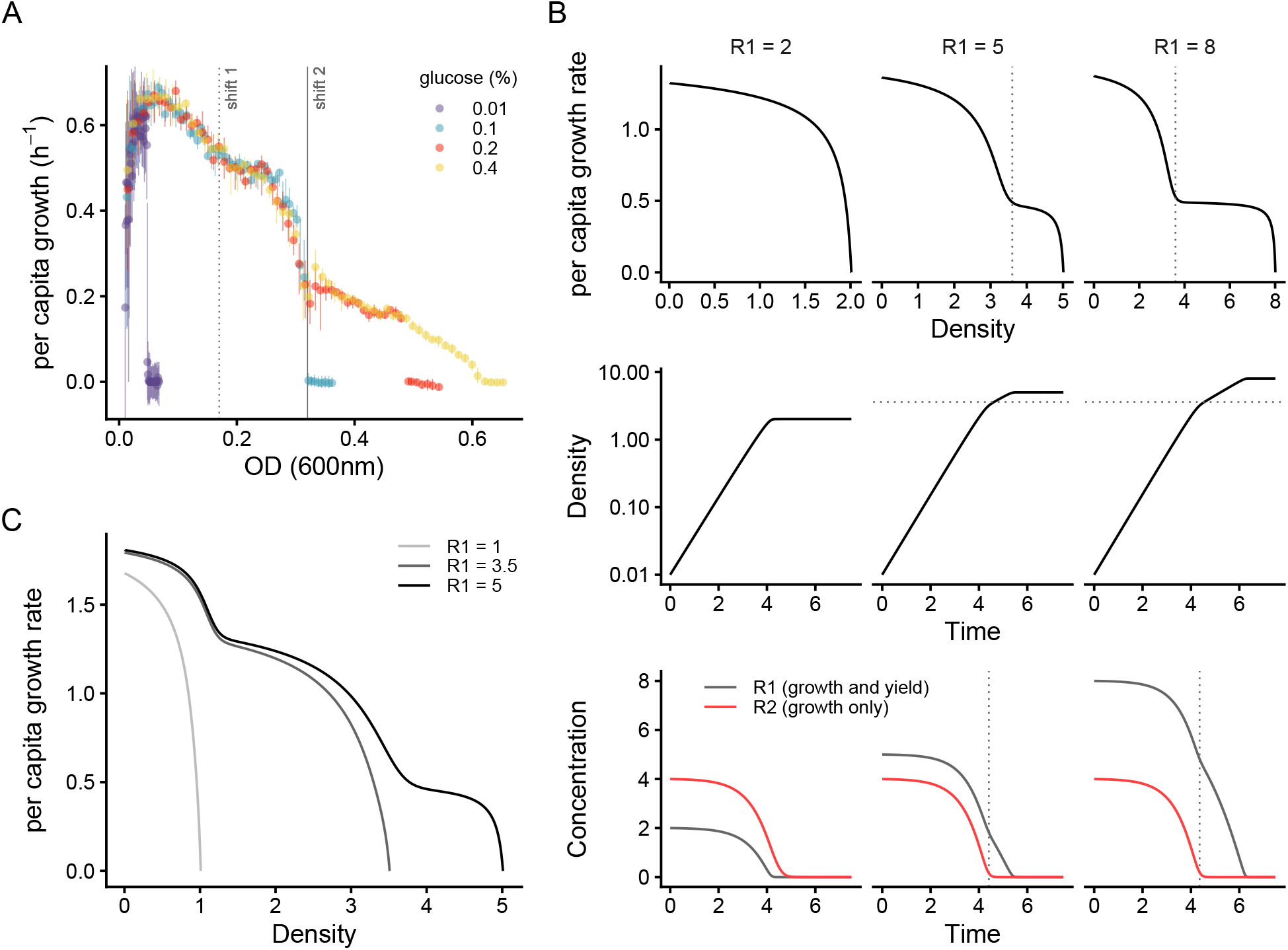
A minimal consumer-resource model reproduces multiphasic patterns of density dependence consistent with observed *E. coli* growth dynamics. **(A)** *E. coli* PGR as a function of optical density as in Fig. 1 but averaged over replicates and across 60 evenly spaced bins for each glucose concentration. Dotted and solid lines denote discrete shifts in growth rate at consistent optical densities (OD_1_=0.17; OD_2_=0.32) despite different initial glucose concentrations. **(B)** Simulations of a model for a single consumer utilizing two resources. Resource 1 limits both growth rate and yield; resource 2 only limits growth rate. Top row: PGR as a function of consumer density when resource 1 is supplied at 0.5x (left column), 1.25x (middle column) or 2x (right column) the concentration of resource 2. Middle row: consumer density as a function of time (y-axis on a log scale) for the corresponding panels in top row. Vertical (top row) and horizontal (middle row) dotted lines denote optical densities where discrete shifts in growth rate are visible. Bottom row: resource concentrations as a function of time for the corresponding panels in the upper rows. Vertical dotted line in bottom row denotes time point at which the discrete shift in growth rate is visible in upper rows. Parameter values are *e*=1.0, *a*_1_=0.6, *k*_1_=0.1, *a*_2_=2.0, *k*_2_=0.5, *α*=1. Initial values are *N* =0.01, *R*_2_=4, and *R*_1_=2 (left column), *R*_1_=2 (middle column), or *R*_1_=8 (right column). **(C)** PGR as a function of consumer density in three simulations of a model for a consumer utilizing three resources (R1 is rate and yield limiting, R2 and R3 are rate limiting only). R1 is supplied at a different initial concentration in each simulation (denoted by line shading), while R2 and R3 are supplied at the same initial concentration in each simulation.

While shifts in growth rate can arise in the presence of multiple substitutable resources (e.g., multiple carbon sources that are consumed sequentially) ^11^, our media only contained a single carbon source, alongside critical macro and micro nutrients (see Methods). Such shifts are also not expected when all resources are purely essential, including under co-limitation ^12^. Instead, the observation that the critical transition between regions of superlinearity and sublinearity occurred at remarkably similar densities—despite different initial glucose concentrations— suggests that these shifts in growth rate may be attributable to the exhaustion of non-essential resources that act primarily as growth enhancers/cofactors (e.g., oxygen or trace metals that become rate limiting well before they become yield limiting, such as Fe, Zn or Mn). To investigate this phenomenon theoretically, we formulated a minimal consumer-resource model for a single consumer utilising two resources, the first of which is both yield and growth rate limiting (e.g., a carbon source) while the second is solely rate limiting:

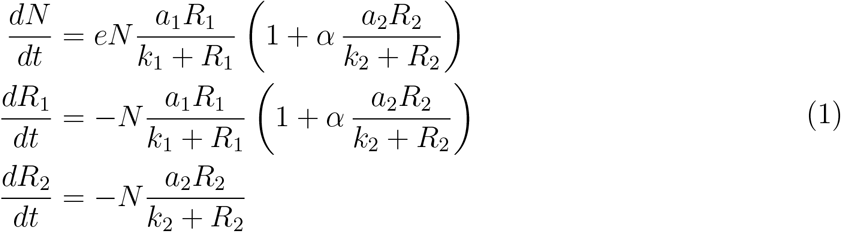

where *N* is the consumer density, *R*_*i*_ is the concentration of resource *i, a*_*i*_ is the maximum uptake rate of resource *i, k*_*i*_ is the half saturation constant on resource *i*, and *e* is the conversion efficiency of resource 1 into consumer biomass. Under this formulation, the first resource (*R*_1_) is essential, mediating both yield and growth rate, while the second resource (*R*_2_) is non-essential but its uptake enhances the consumer’s growth rate (the strength of which is regulated by *α*).

We simulated this model under a range of different resource uptake rates, half saturation constants, and initial resource concentrations for the two resources in batch culture (see Methods). As exemplified in Fig. 2B, the dominant pattern that emerges is consistent with our experimental observations, whereby an excess of the yield-limiting resource results in multi-phasic density dependence: growth initially declines superlinearly until the exhaustion of non-essential *R*_2_ (grey dotted line in bottom row of Fig. 2B) precipitates a sublinear transition to a new phase of superlinear density dependence. In addition, as with the experimental data, these transitions occur at the same density even under substantial differences in the concentration of the yield limiting resource. Note, the abruptness of the transition and the prominence of the two phases varies by growth parameters and initial resource concentrations (Figs. S1 to S4).

A constraint of this basic two resource model is that it only produces a single region of sublinearity, and yet our experimental data was characterised by two distinct sublinear regions. (Note that each glucose concentration is characterised by an artefactural sublinear region at PGR *≈* 0 in Fig. 2A, which emerges from averaging growth rate at carrying capacity across multiple replicates, each with small variation in inoculum densities.) To see if we could reproduce dynamics with two distinct shifts, we subsequently simulated an extended model with an additional rate limiting resource (see Methods), also across a range of parametrisations and initial concentrations. This model produced dynamics that were closely aligned with the experimental data, with the initial concentration of the yield limiting resource strongly determining whether none, one, or two sublinear shifts are visible (Fig. 2C, Extended Data Fig. 1).

Finally, to evaluate the potential ubiquity of multi-phasic patterns of density dependence under multiple resource limitation, we also explored alternative formulations of the basic model, including a more canonical consumer-resource model for additive, substitutable resources (see Methods). These model variations produced similar multi-phasic patterns of density dependence to those presented in Fig. 2 (Figs. S5 and S6). While the assumption of substitutable resources is almost certainly incompatible with *E. coli* growth on glucose in M9 media, the substitutable-resources model reasonably captures the basic consumer-resource dynamics of many macro-organisms (see also Ref. 13 for similar observations with a differently formulated model).

Our findings provide an alternative explanation for recent empirical observations of sublinearity^4,5^. In particular, we suggest that the reported sublinearity in *E. coli* in Ref. 5 is driven by shifts in the identity of the growth rate-limiting resource, which leads to the signature transition between regions of super- and sublinearity. Although sequential shifts in resource limitation can lead to almost linear density dependence at low density (e.g., Fig. S1), we posit that density dependence will most typically be characterised by superlinearity at low (e.g., when colonising an unexploited environment) and high density (e.g., around equilibrium). The magnitude of any sublinear transitions in between would then depend on the availability of multiple growth-limiting resources and how rapidly these become exhausted (e.g., as influenced by their uptake rate and half-saturation constant).

Although our empirical observations derive from simple experiments with a model bacterium, the underlying mechanisms are likely to play out in more complex macro-organismal systems. This is readily illustrated by the emergence of sublinearity at intermediate densities out of standard substitutable-resource models (Fig. S6 and Ref. 13), which are commonly used in mechanistic modelling of animal populations^14^. More generally, we have likely only scratched the surface of resource-based interactions that can yield emergent sublinearity^12,15,16^. Indeed, Abrams (2022) ^17^ presents a model of adaptive foraging for two resources based on resource abundance that yields very similar patterns of density dependence to those described here (i.e., superlinear density dependence at low and high density joined by a region of sublinearity at intermediate densities).

Despite providing a parsimonious resource-based explanation for multiphasic patterns of density dependence, it would be premature to conclude that sequential resource use is the primary mechanism underlying patterns of sublinearity observed in nature. Testing this hypothesis using observational time series presents a formidable challenge, not least because it requires sufficiently high-resolution data to reliably fit models characterised by multiple inflections in density-dependent per capita growth. Moreover, significant caution is needed when drawing inference on patterns of density dependence from coarse time-series data (particularly in macro-organisms), where measurement error, low temporal resolution, and delayed effects can generate artefactual sublinearity^7–9,13^. As such, progress towards a more mechanistic understanding of density dependence will likely depend on carefully designed studies that integrate manipulative experiments alongside traditional time-series analyses.

### Online Methods

#### Growth assays

Prior to conducting the experimental growth assays, three overnight cultures of *E. coli* strain MG1655 were grown in M9 + 0.1% glucose for 18 hours, before pelleting 1mL of culture from each overnight, and resuspending the pellet in 200*µ*L M9 (0% glucose). Pelleting and resuspension was implemented to remove residual glucose prior to inoculation in the experimental plate. 2*µ*L of the resuspension from each overnight was then inoculated in triplicate into 178*µ*L of M9 across a glucose gradient (0.01%, 0.1%, 0.2%, 0.4%) in a 96 well plate, for a total of nine replicates per glucose concentration (three biological replicates x three technical replicates). 180*µ*L of media at each glucose concentration was added to each of four wells without inocula to serve as blanks. Optical density (OD) was read every three minutes in a Biotek Epoch 2 plate reader for 24 hours. The 0.01% glucose treatment was repeated a further two times on independent plates, for a total of 27 replicates.

### Analysis of growth data

Raw data from the plate reader was imported into R (v4.5.0) via the *grow96* package (https://github.com/JanEngelstaedter/grow96). OD measurements were blanked by subtracting OD readings at each time point in the blank wells from experimental wells of the same initial glucose concentration. Per capita growth rate, *g*_*t*_, in each well at each time point was calculated using the central difference approximation based on the difference in the natural log of OD and time elapsed across each successive interval:

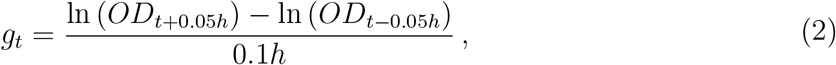

which gives estimates in units of inverse hours. Plots of the resulting density dependence were therefore based on a comparison of *g*_*t*_ against *OD*_*t*_.

Owing to spuriously high and low estimates of per capita growth when densities were close to the detection limit of the plate reader, the first hour of readings was ignored for all wells prior to analysing the data. In addition, we excluded all measurements taken more than *>*∼1h post saturation. This time cut-off varied by initial glucose concentration as follows: 0.01% = 4h, 0.1% = 7.5h, 0.2% = 10h, 0.4% = 15h.

### Model simulations

All simulations were run in R (v4.5.0) using the lsoda solver from the deSolve package (v1.4)^18^. Simulations were initiated with consumer density at 0.01, and integrated over 20 units of time, with solutions reported every 0.01 time-steps. See main text for model formulations and parametrization.

## Code availability

R code to reproduce simulations and analysis will be deposited in a publicly available repository.

## Acknowledgements

We thank Joseph Wenck and Andrew Tuck for support in conducting the bacterial growth assays. ADL is supported by Australian Research Council grants DP220103350 and DE230100373.

## Extended Data

**Extended Data Figure 1:**
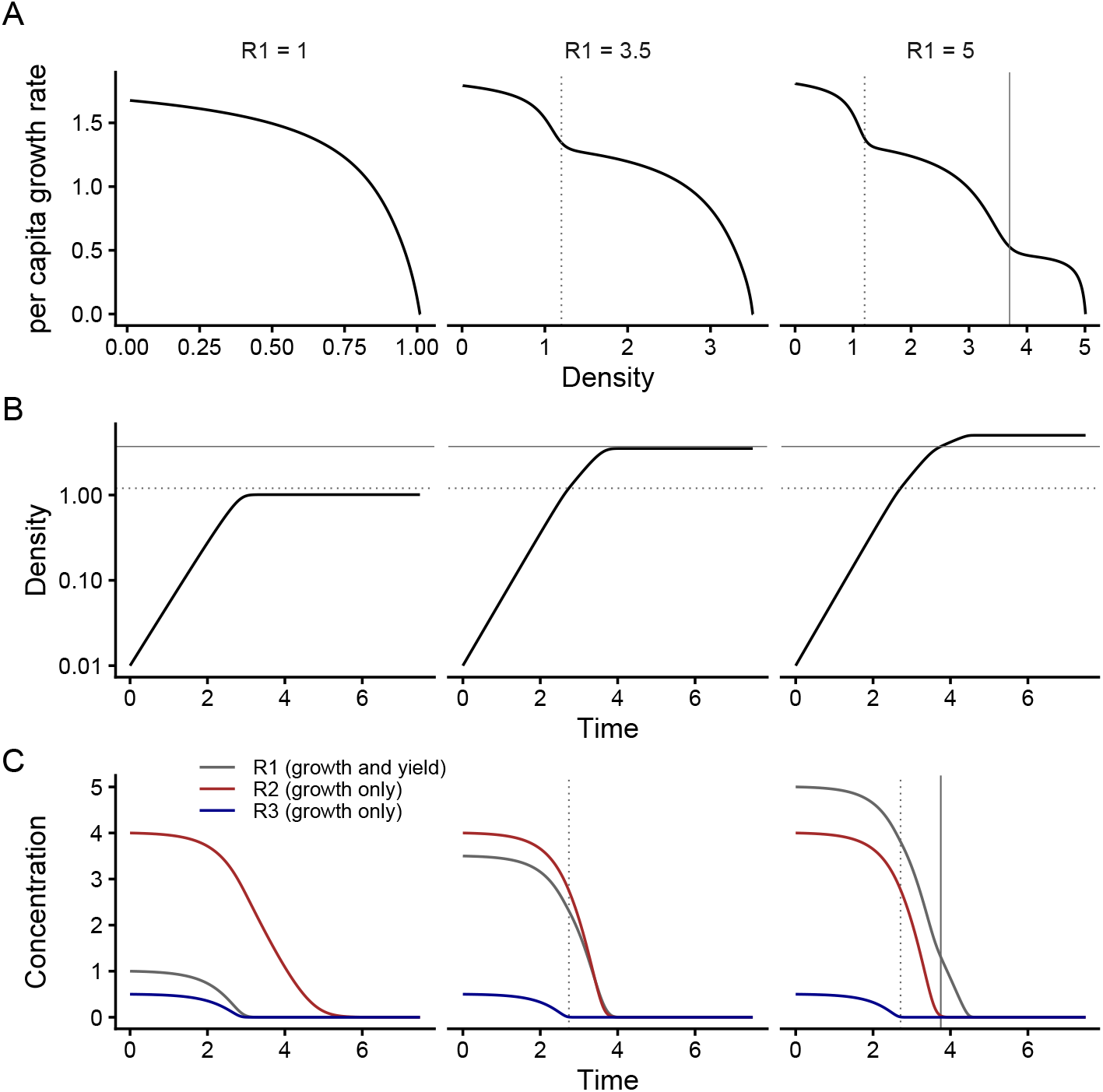
Simulations of a model for a single consumer utilizing three resources. Resource 1 limits both growth rate and yield; resources 2 & 3 only limit growth rate. Top row: PGR as a function of consumer density when resource 1 is supplied at low (left column), medium (middle column) and high (right column) concentration. Middle row: consumer density as a function of time (y-axis on a log scale) for the corresponding panels in top row. Vertical (top row) and horizontal (middle row) dotted lines denote optical densities where discrete shifts in growth rate are visible. Bottom row: resource concentrations as a function of time for the corresponding panels in the upper rows. Vertical dotted line in bottom row denotes time point at which the discrete shift in growth rate is visible in upper rows. Parameter values are *e*=1.0, *a*_1_=0.5, *k*_1_=0.1, *a*_2_=2.0, *k*_2_=0.5, *α*_2_=1, *a*_3_=1.0, *k*_3_=0.05, *α*_3_=1. Initial values are *N* =0.01, *R*_2_=4, *R*_2_=0.5, and *R*_1_=1 (left column), *R*_1_=3.5 (middle column), or *R*_1_=5 (right column).

## Supplementary Information

Additional supporting information may be found in the online version of this article.

## Supplementary Methods

### Three-resource model formulation

The three-resource model discussed in the main text took the following form:

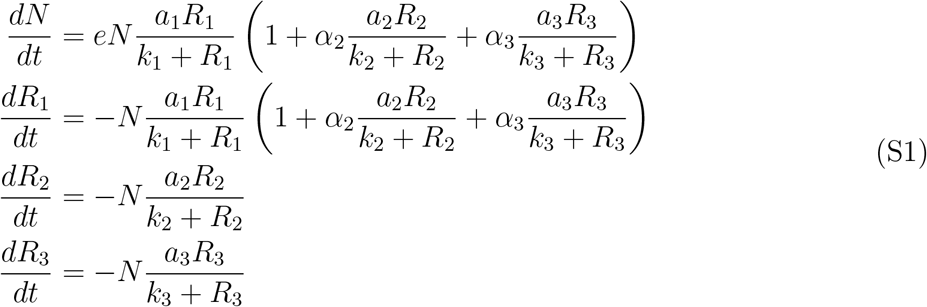

where R1 is rate and yield limiting, while R2 & R3 are rate limiting only.

### Alternative two-resource model formulations

Alongside the core model presented in the main text we also simulated dynamics under two alternative model formulations.

The first alternative model is of the following form:

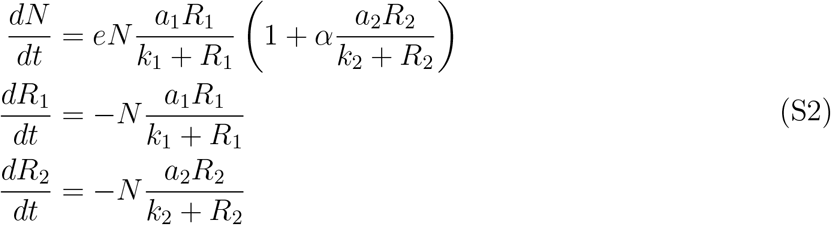

Under this formulation, *R*_2_ still affects the growth of the consumer but does not affect the uptake of *R*_1_. Example simulation output shown in Fig S5.

The second alternative model is a classical consumer-resource model for two substitutable resources:

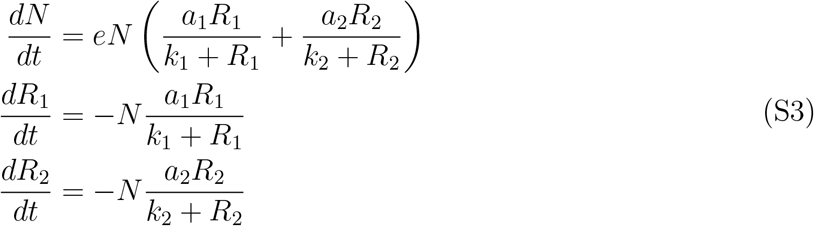

Example simulation output shown in Fig S6.

**Figure S1:**
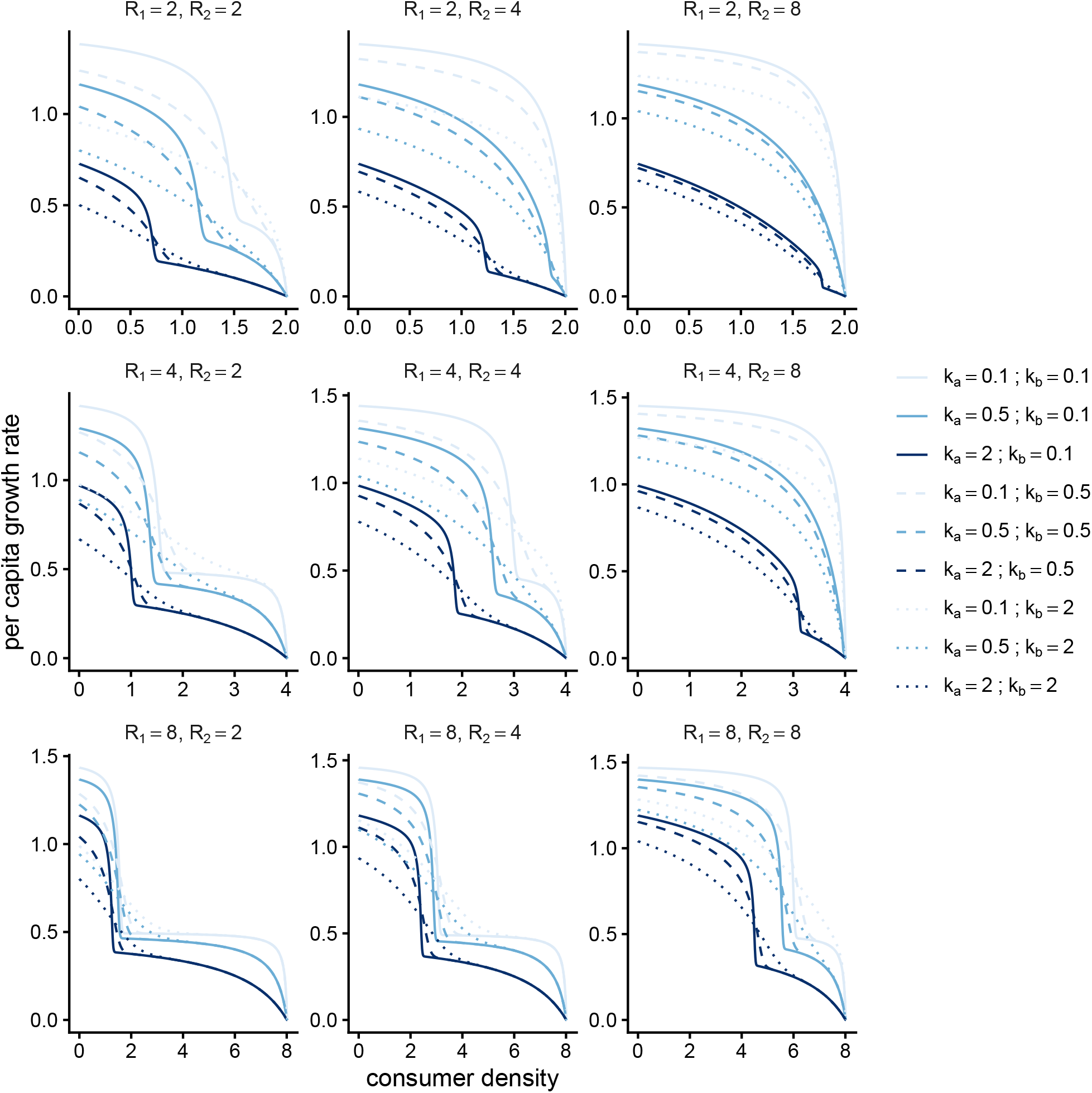
PGR as a function of consumer density in three simulations of a model for a single consumer utilizing two resources. Resource 1 limits both growth rate and yield; resource 2 only limits growth rate. Parameter values are *e*=1.0, *a*_1_=0.5, *a*_2_=2.0, *k*_*i*_=see legend. Individual panels denote initial resource values.

**Figure S2:**
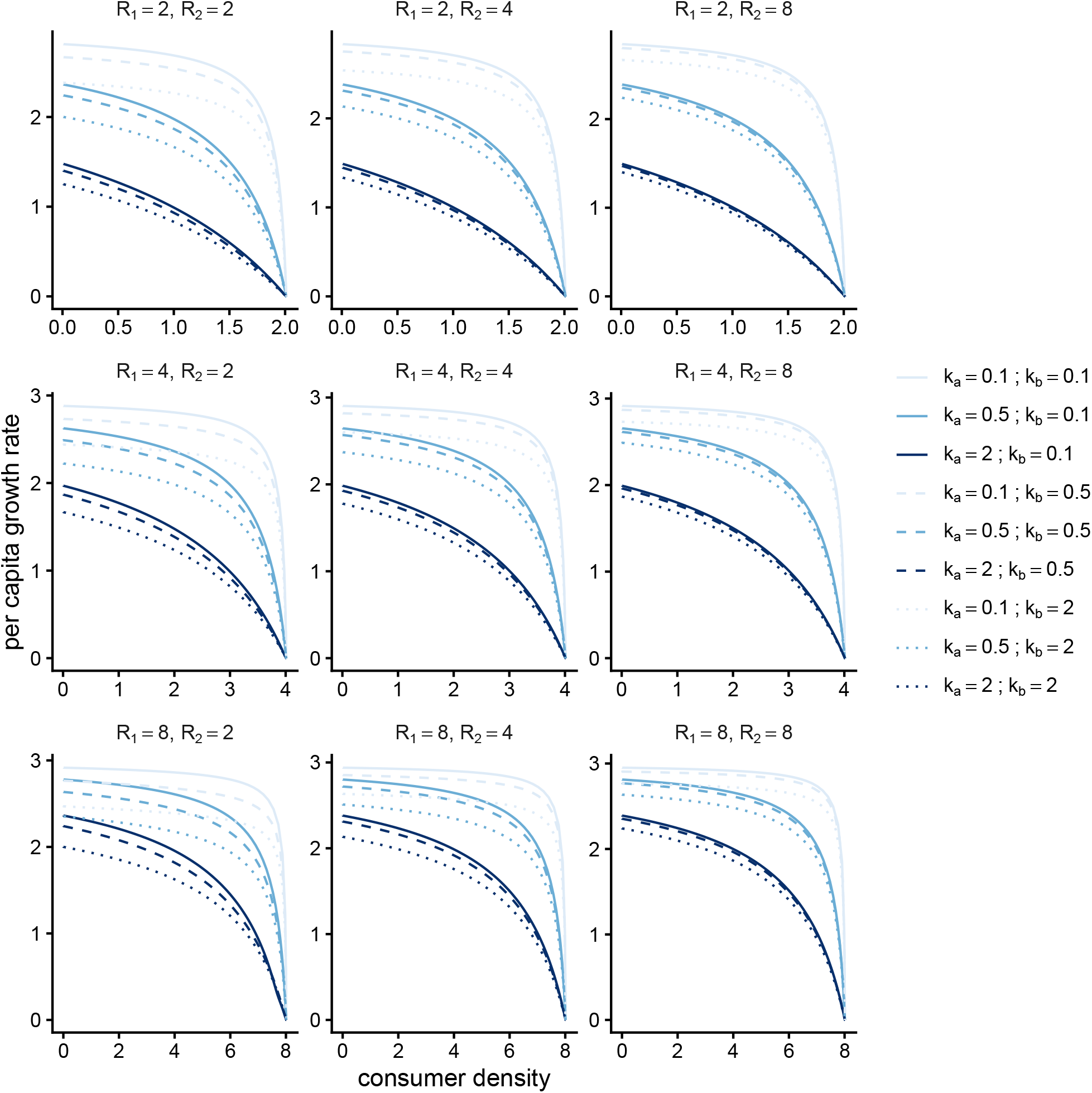
PGR as a function of consumer density in three simulations of a model for a single consumer utilizing two resources. Resource 1 limits both growth rate and yield; resource 2 only limits growth rate. Parameter values are *e*=1.0, *a*_1_=2, *a*_2_=0.5, *k*_*i*_=see legend. Individual panels denote initial resource values.

**Figure S3:**
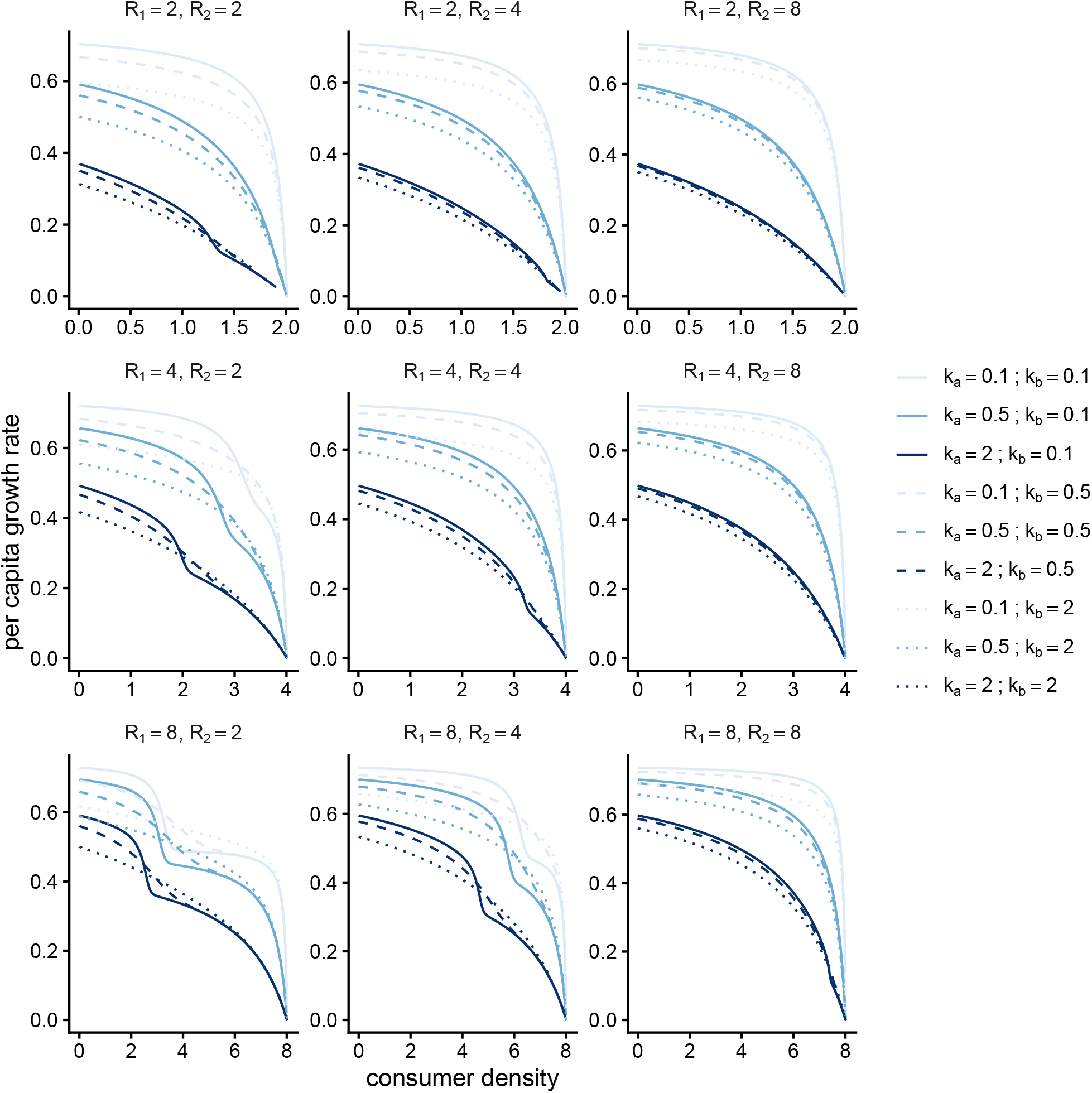
PGR as a function of consumer density in three simulations of a model for a single consumer utilizing two resources. Resource 1 limits both growth rate and yield; resource 2 only limits growth rate. Parameter values are *e*=1.0, *a*_1_=0.5, *a*_2_=0.5, *k*_*i*_=see legend. Individual panels denote initial resource values.

**Figure S4:**
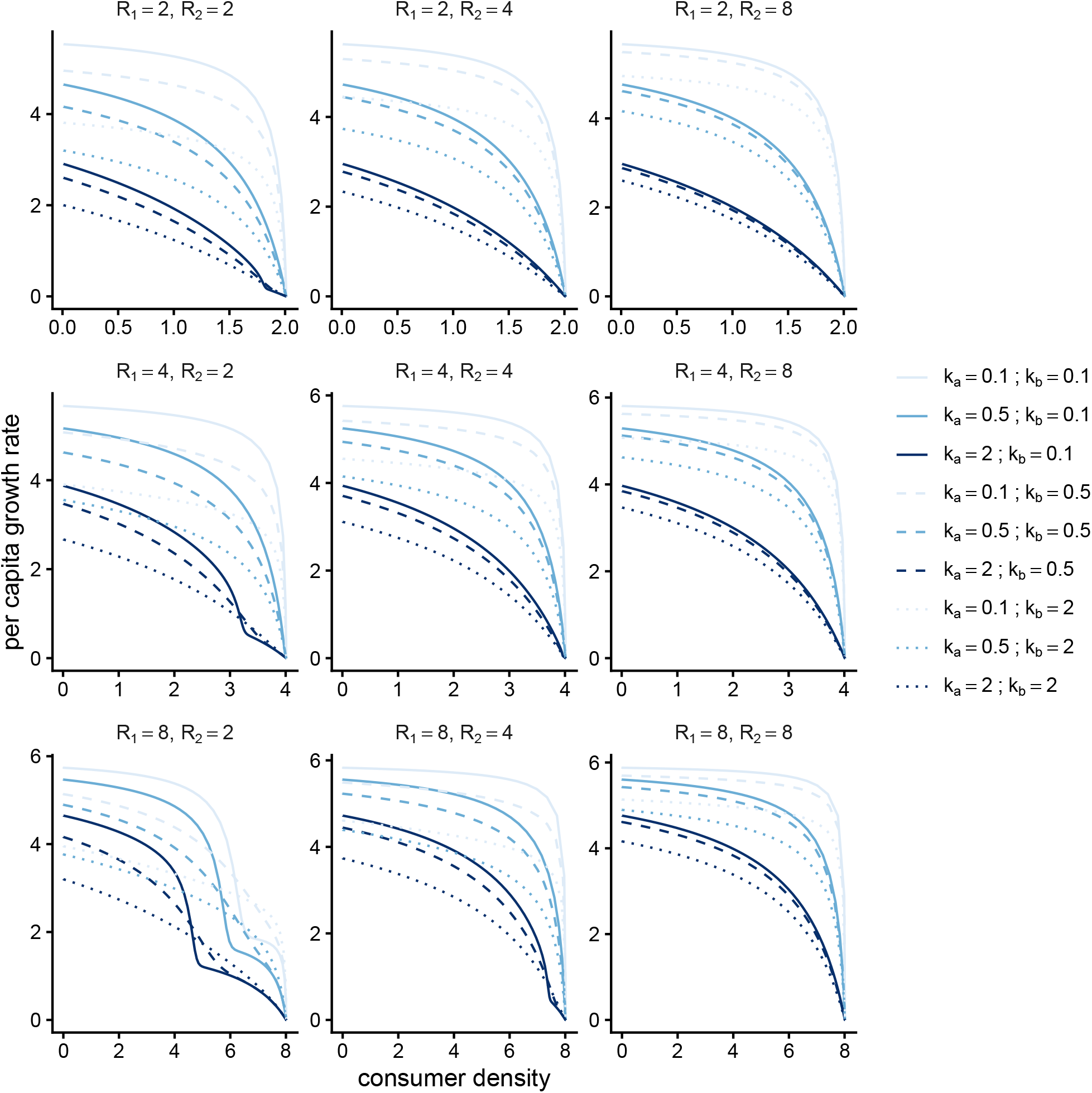
PGR as a function of consumer density in three simulations of a model for a single consumer utilizing two resources. Resource 1 limits both growth rate and yield; resource 2 only limits growth rate. Parameter values are *e*=1.0, *a*_1_=2, *a*_2_=2.0, *k*_*i*_=see legend. Individual panels denote initial resource values.

**Figure S5:**
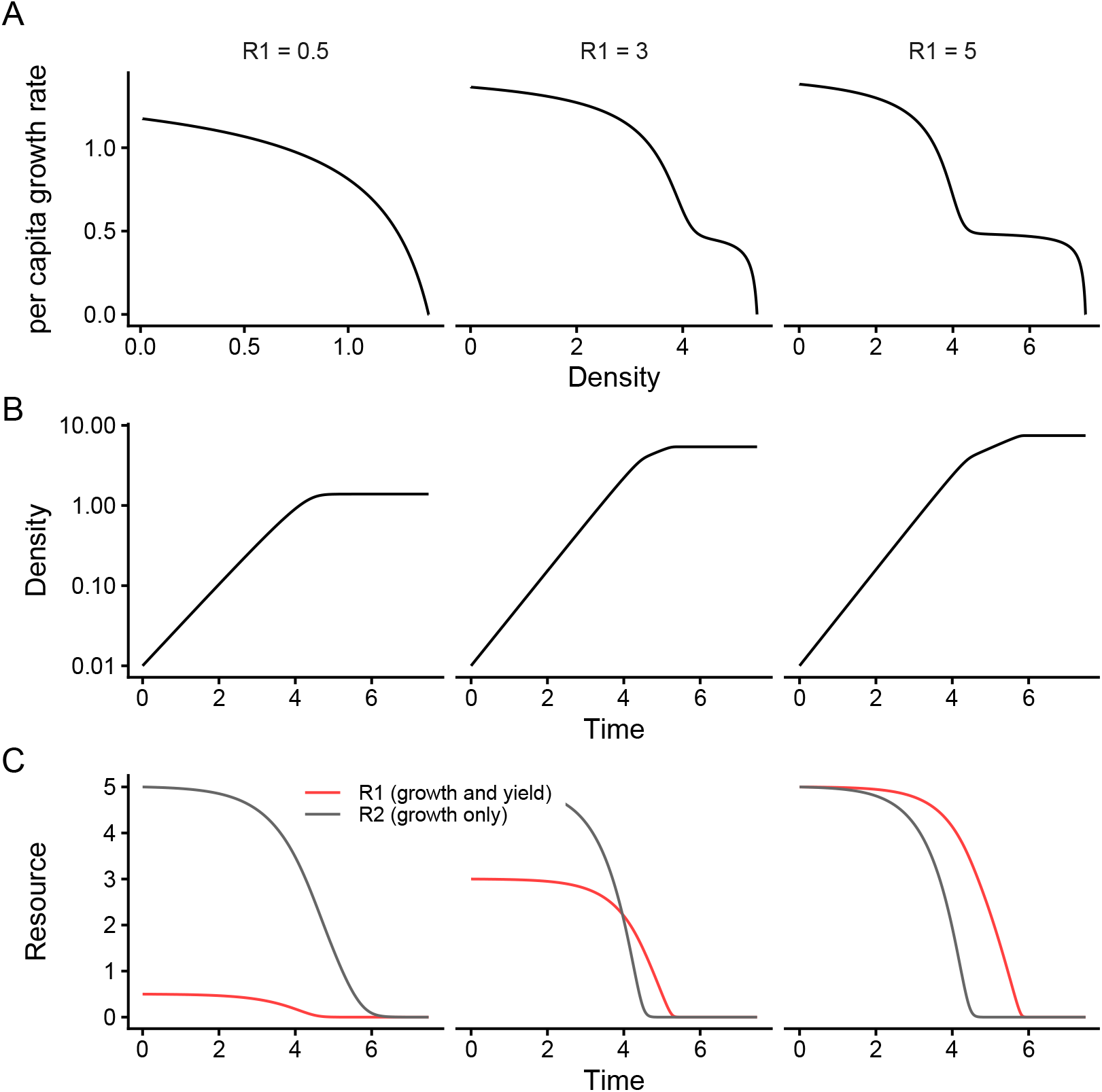
Simulations of a model for a single consumer utilizing two resources (based on Eq S2), where resource 1 limits both growth rate and yield, while resource 2 only limits growth rate. Top row: PGR as a function of consumer density when resource 1 is supplied at low (left column), medium (middle column) and high (right column) concentration. Middle row: consumer density as a function of time (y-axis on a log scale) for the corresponding panels in top row. Vertical (top row) and horizontal (middle row) dotted lines denote optical densities where discrete shifts in growth rate are visible. Bottom row: resource concentrations as a function of time for the corresponding panels in the upper rows. Vertical dotted line in bottom row denotes time point at which the discrete shift in growth rate is visible in upper rows. Parameter values are *e*=1.0, *a*_1_=0.5, *k*_1_=0.1, *a*_2_=2.0, *k*_2_=0.5, *a*_2_=1.0, *k*_2_=0.05. Initial values are *N* =0.01, *R*_2_=5, and *R*_1_=0.5 (left column), *R*_1_=3 (middle column), or *R*_1_=5 (right column).

**Figure S6:**
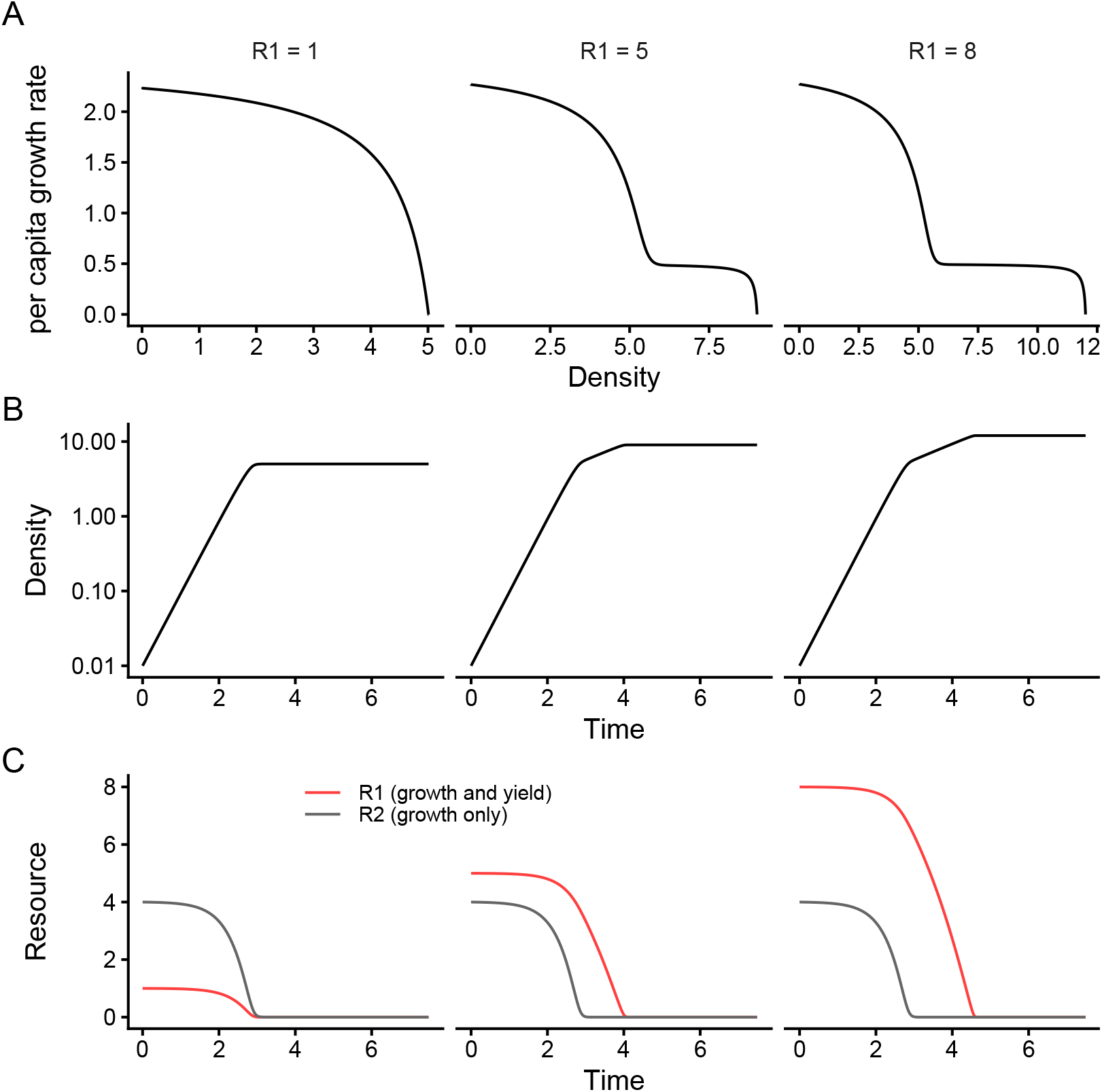
Simulations of a model for a single consumer utilizing two resources (based on Eq S3), where resource 1 limits both growth rate and yield, while resource 2 only limits growth rate. Top row: per capita growth rate (PGR) as a function of consumer density when resource 1 is supplied at low (left column), medium (middle column) and high (right column) concentration. Middle row: Consumer density as a function of time (y-axis on a log scale) for the corresponding panels in top row. Vertical (top row) and horizontal (middle row) dotted lines denote optical densities where discrete shifts in growth rate are visible. Bottom row: Resource concentrations as a function of time for the corresponding panels in the upper rows. Vertical dotted line in bottom row denotes time point at which the discrete shift in growth rate is visible in upper rows. Parameter values are *e*=1.0, *a*_1_=0.5, *k*_1_=0.1, *a*_2_=2.0, *k*_2_=0.5, *a*_2_=1.0, *k*_2_=0.05. Initial values are *N* =0.01, *R*_2_=4, and *R*_1_=1 (left column), *R*_1_=5 (middle column), or *R*_1_=8 (right column).

